# Utilizing a pan-transcriptome reveals genotype-specific responses to iron deficiency in *Sorghum bicolor*

**DOI:** 10.1101/2024.12.17.626333

**Authors:** Chinaza Davies Nnamdi, Kittikun Songsomboon, Azita Alasvand Zarasvand, William G. Voelker, Elizabeth A. Cooper

## Abstract

**Introduction:** Sweet sorghum genotypes are distinguished from non-sweet types by their ability to accumulate high concentrations of sugar in their stems, which makes them valuable for both bioenergy production and as an alternative source of sugar. Previous work has reported numerous gene copy number differences in metal metabolism and iron transport genes between the sweet and non-sweet genotypes. The objective of this study was to investigate which genes are differentially expressed by the sweet and non-sweet genotypes under varying iron conditions.

**Methods:** Three sweet and three non-sweet sorghum genotypes were grown hydroponically under two conditions (control and low iron), and root tissue from the plants was harvested for RNA extraction and sequencing after 9 days. To obtain a more complete picture of diverse iron-response mechanisms in sorghum, we constructed a pan-transcriptome from 10 founder parents of the Carbon-Partitioning Nested Association Mapping population.

**Results:** Differential expression analysis using the pan-transcriptome revealed 209 genes that responded differently to iron stress in the sweet and non-sweet types, which was more than twice as many genes found using only a single reference genome. Enrichment analysis based on age of duplication event showed that duplicate gene copies that are differentially expressed due to genotype effect appear to be specific to sorghum while most duplicate gene copies that are differentially expressed due to iron treatment appear to be more evolutionary conserved across plants.

**Discussion:** This study highlights the importance of the non-core genome and the utility of the pan-transcriptome in sorghum. It also revealed that different sorghum genotypes respond to iron stress differently.

## 1 Introduction

*Sorghum bicolor* (L.) Moench, commonly known as sorghum, is an important source of food, feed, and biofuel. There are four major agronomic sorghum types: sweet, grain, forage, and cellulosic (Brenton et al., 2016; Kumar et al., 2022). The sweet genotypes are characterized by their potential to accumulate high concentrations of sugar in their stems which makes them useful for both bioenergy production and as an alternative source of sugar (Murray et al., 2009; Burks et al., 2015).

Previous work has reported numerous variations in metal metabolism and iron transport genes between the sweet and non-sweet sorghum genotypes. For instance, a comparative transcriptomic study of a sweet type (Rio) and a non-sweet type (PR22) noted that iron-transport related genes were differentially expressed between the two types over the course of plant development (Cooper et al. (2019). Songsomboon et al. (2021) reported that metal stress-related genes were enriched for population-specific deletions and Voelker et al. (2023) showed that metal transporters were frequently impacted by structural variants specifically in sweet types. A Genome-Wide Association Study (GWAS) utilizing a panel that comprised sweet and non-sweet types identified a sorghum-specific copy of a putative vacuolar iron transporter (*VIT*) as a top candidate gene for sugar content (Brenton et al., 2016; Brenton et al., 2020).

While the precise function and role of the genomic variation in iron transporters in sweet sorghums remains unknown, these studies strongly suggest that differences in iron metabolism could be linked to stem sugar accumulation in sorghum. Iron is an essential micronutrient in plants; it plays a role in sugar metabolism by being a crucial constituent of several enzymes and components involved in processes related to sugar synthesis, transport and storage (Marschner, 2011; Briat et al., 2015; Rout and Sahoo, 2015; Ning et al., 2023). For example, iron plays a role in the synthesis of chlorophyll, and it is a component of the major complexes of the photosynthetic apparatus (Marschner, 2011; Briat et al., 2015). Efficient photosynthesis is important to produce sugar, which is fundamental for plant growth and development (Marschner, 2011). Iron is also a component of aconitase, a key enzyme in the citric acid cycle and iron is a component of cytochromes, an important enzyme in electron transport and respiration (Marschner, 2011; Rout and Sahoo, 2015).

Given all the previous work showing variability of metal and iron transport genes between the sweet and non-sweet types and the potential link between improved iron metabolism and increased stem sugars, the goal of this study was to investigate whether or not the different types actually respond differently to iron stress at the genetic level. This was conducted using differential expression analyses of the different sorghum types under varying iron conditions. Given that the iron transport gene reported in the earlier GWAS was shown to be the result of a tandem duplication specific to sorghum, this study also investigated the evolutionary origins of differentially expressed genes in order to determine if other sorghum-specific gene copies might be driving differences between the two types.

When considering the effects of newer genes, it is important to note that a single reference genome or transcriptome from only one individual does not accurately reflect the diversity of gene content that can occur within a species. To overcome these limitations, researchers have recently been constructing pan-genomes and pan-transcriptomes and using them in their studies (Golicz et al., 2016; Della Coletta et al., 2021; Kong et al., 2022; Zhang et al., 2022; Yan et al., 2023). In a striking example of this, Ma et al. (2019) constructed a pan-transcriptome for barley from 63 different genotypes (that comprised cultivated and wild types) and found that 38.2% of the transcripts in the pan-transcriptome were not present in the original reference genome. This study also reported the prevalence of genes associated with responses to different stresses and stimuli among the novel transcripts.

In this study, we performed differential expression analyses using both a pan-transcriptome and a more traditional single reference genome approach. We constructed our pan-transcriptome from 10 (Grassl, PI329311, PI506069, PI510757, Chinese Amber, Rio_NAM, Leoti, PI229841, PI297155, PI655972) of the 12 founder parents of the Carbon-Partitioning Nested Association Mapping (CP-NAM) population (Boatwright et al., 2021). These lines are very diverse and represent the four major sorghum types (sweet, grain, forage, and cellulosic) and the five sorghum races (bicolor, caudatum, durra, guinea, and kafir) (Boatwright et al., 2021; Kumar et al., 2022; Voelker et al., 2023). We compared our results from the pan-transcriptome analysis to the results we obtained when aligning reads to the representative sweet sorghum (‘Rio’) genome to evaluate the advantage of the pan-transcriptome.

This study revealed that genes are differentially expressed in different sorghum genotypes in response to iron stress and showcased the utility of the pan-transcriptome in the identification of more differentially expressed genes. This study also showed that duplicate gene copies that are differentially expressed due to genotype effect appear to be specific to sorghum while most genes that are differentially expressed due to only iron treatment are more evolutionary conserved across plants.

## 2 Materials and Methods

### 2.1 Sorghum genotypes and growth conditions

Three sweet (Grassl, Leoti, Rio_NAM) and three non-sweet (PI506069, PI655972, PI297155) sorghum genotypes were utilized in this study. Seeds from all genotypes were obtained from the Germplasm Resources Information Network; they were treated with Captan fungicide and germinated in Petri dishes with wet filter papers for five days at room temperature. Healthy seedlings were transplanted to grow hydroponically in 1/5 Hoagland solution at pH 5.5 under T5 fluorescent 20,000 lumens 16-hour light and aerated 24 hours for five days. After five days, genotypes were subjected to two iron conditions (control - 100 μM Fe and low iron - 0 μM Fe). Iron treatments were applied by full-strength Hoagland solution (4.0 mM Ca(NO_3_)_2_, 2.0 mM MgSO_4_, 1.0 mM NH_4_H_2_PO_4_, 6.0 mM KNO_3_, 3.0 μM H_3_BO_3_, 0.5 μM MnSO_4_, 0.4 μM ZnSO_4_, 0.2 μM CuSO_4_, and 0.05 μM H_2_MoO_4_). Each genotype-by-condition treatment had 3 biological replicates and the experimental design was a randomized complete block design. There was no replicate within a block and each block contained one of each of the 6 different genotypes. The pH was adjusted daily to 5.5. All the hydroponic systems were set up in a growth chamber at 32°C both 16 h day and 8 h night, with a relative humidity of 55%, and light intensity of 2,700 lumens from T5 fluorescence supplement light. After 9 days of iron treatment, bulk roots from each individual (∼100 mg) were harvested and stabilized by RNAlater at 8°C for 24 hours. After that, excessive RNAlater solution was drained, and RNA-stabilized root tissues were kept at -80°C for further RNA extraction.

### 2.2 mRNA extraction, RNA seq library preparation, and RNAseq of sorghum

RNA-stabilized and frozen sorghum root tissue from each individual was ground and the RNA was extracted with an RNeasy mini kit (Qiagen) according to the manufacturer’s instructions. Total RNA concentrations were measured using the Qubit® Broad Range RNA kit on a Qubit fluorometer (Q32857), and the purity of RNA was determined by the Nanodrop spectrophotometer 2000. RNAseq library preparation and sequencing were conducted by David H. Murdock Research Institute. Due to the low quantity of high-quality RNAseq reads obtained from some replicates, only five sorghum genotypes (Grassl, Leoti, Rio_NAM, PI506069, PI655972) and certain replicates were included in further analysis (Supplementary Table 1).

### 2.3 Pan-transcriptome construction

The cDNA sequence files from each of the 10 founder parents of CP-NAMS were obtained from SorghumBase (https://www.sorghumbase.org/) (Gladman et al., 2022). Voelker et al. (2023) previously constructed *de novo* genome assemblies and annotations for these 10 lines and provided a presence/absence variation (PAV) file that detailed which of the lines had a gene for each of the annotated 62,044 loci identified in the pan-genome. A Python script was written to get the longest sequence for each locus to construct the complete pan-transcriptome FASTA file (Supplementary Figure 1 and Supplementary File 1).

### 2.4 Differential expression analysis

Raw RNA-seq reads were trimmed and cleaned using Trimmomatic v0.38 with the following parameters: PE-phred33 ILLUMINACLIP:TruSeq3-PE.fa:2:30:10 LEADING:3 TRAILING:20 SLIDINGWINDOW:4:15 MINLEN:36 (Bolger et al., 2014). Alignment of the trimmed reads to the Rio reference genome was done using STAR v2.7.9a with the options –alignEndsType Local --alignIntronMax 0 --outSAMtype BAM SortedByCoordinate --outSAMunmapped Within --outSAMattributes Standard (Dobin et al., 2012). STAR was also used in aligning the trimmed reads to the pan-transcriptome with options --alignEndsType EndToEnd --alignIntronMax 1 --outSAMtype BAM SortedByCoordinate --outSAMunmapped Within --outSAMattributes Standard. STAR is a splice-aware aligner; to turn off its detection of unannotated splice junctions, alignIntronMax was set to 1. To turn off its detection of annotated splice junctions, genome indexing was done without any GTF or GFF annotation file. Since the pan-transcriptome has no intron regions, the option --alignEndsType EndToEnd was used to prevent soft-trimming or soft-clipping of read characters during alignment.

Read counts were obtained using mmquant; this tool has an advantage over similar read count tools by not automatically discarding reads that map to multiple locations in the reference sequence, which would be potentially highly problematic when mapping to a pan-transcriptome that contains many similar duplicate gene copies. Rather than discarding these multi-mapping reads, it creates new “hybrid” gene identifiers and assigns the multi-mapping read counts to this new ID (Zytnicki, 2017). To use mmquant with our pan-transcriptome, a Python script was written to make a pseudo-GTF file for the pan-transcriptome (Supplementary File 2).

Differential expression analysis was done using the glmmSeq package in R (Lewis et al., 2022), which modeled the data using a negative binomial mixed effects model. Treatments (Control vs Low Iron) and Type (Sweet vs Non-sweet) were treated as fixed effects while Genotype was treated as a random effect to account for genotype relatedness:

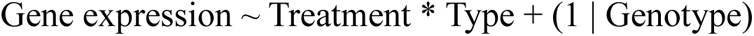

We also performed randomized permutation tests to validate the differences between sweet and non-sweet types (see Supplementary Material for details).

To effectively compare the differentially expressed genes (DEGs) from the analysis using the Rio reference genome and the analysis using the pan-transcriptome, we used Orthofinder v2.5.4 on the protein sequences from the Rio reference genome and the lines that make up the pan-transcriptome to group genes into orthogroups that can be inferred to have descended from a single common ancestor (Emms and Kelly, 2019). Afterward, Python and R scripts were written to parse the result and generate Venn diagrams that showed overlapping orthogroups of the DEGs from both analyses for each of the fixed effects in the model.

To determine the functional descriptions of DEGs, a Python script was written to extract this information from the National Center for Biotechnology Information (NCBI). A BLAST search was done on genes that had no description on NCBI or that were described as a hypothetical protein or as an uncharacterized protein. GNU Parallel 20210222 was used during the BLAST search to accelerate the process (Tange, 2021). The top 10 results of the BLAST search were kept. This information is available in Supplementary File 3.

### 2.5 Gene Ontology enrichment analysis

The annotation file for the Rio reference genome was obtained from (Cooper et al., 2019), and an R script was written to parse this file and extract the Gene Ontology (GO) terms.

InterProScan v5.60-92.0 was used in getting the GO annotation of proteins of the 10 founder parents of CP-NAMS (Jones et al., 2014). A Python script was written to parse the result generated by InterProScan and annotate the pan-transcriptome.

Gene ontology (GO) enrichment analysis for molecular function was performed using the classic algorithm and Fisher’s exact test with a p-value cut-off of 0.05; this was implemented in R using the topGO package (Alexa and Rahnenfuhrer, 2024). GO-Figure, a Python package, was used to calculate the pairwise semantic similarities of GO terms (Reijnders and Waterhouse, 2021). GO-Figure does this using Lin’s formula (Lin, 1998), and a cutoff threshold of 0.7 was set. This means that terms with pairwise semantic similarity scores higher than or equal to 0.7 were grouped into one and a representative term was selected from the group. R was used in plotting representative terms in two-dimensional semantic similarity space; terms with more functional similarities are closer to each other on the scatterplot.

### 2.6 Enrichment analysis based on evolutionary nodes annotation

Orthofinder v2.5.4 was used in performing orthology analysis on selected plant species: *Amborella trichopoda, Ananas comosus, Aquilegia coerulea, Arabidopsis thaliana, Asparagus officinalis, Brachypodium distachyon, Medicago truncatula, Mimulus guttatus, Miscanthus sinensis, Oryza sativa, Panicum hallii, Populus trichocarpa, Setaria italica, Sphagnum fallax, Spinacia oleracea,* and *Zea mays*. Protein sequences from each plant species were obtained from Phytozome (Goodstein et al., 2012) and used as input for Orthofinder.

The results from the analysis using Orthofinder comprise information on orthogroups, orthologs, rooted gene trees for all orthogroups, rooted species trees, and gene duplication events (Emms and Kelly, 2019). Python scripts were written to parse these results and annotate genes that are duplicate gene copies based on their oldest species tree node label which represents their age of duplication event. Genes that were single-copy orthologues and those that were unassigned to any orthogroup were annotated as ‘Single Copy Orthologues’ and ‘No Orthogroup’, respectively. The annotation file generated was used in an enrichment analysis to determine nodes enriched among the DEGs. The enrichment analysis was done using the R package clusterProfiler v4.0 (Wu et al., 2021). clusterProfiler calculated a p-value using hypergeometric distribution, and multiple testing was corrected using the Benjamini-Hochberg method to produce an adjusted p-value (Benjamini and Hochberg, 1995).

## 3 Results

### 3.1 Read alignment and count

After trimming, the average number of surviving reads across the samples was about 16 million (range 10.5-21 million) (Supplementary Table 2). Across the samples, a lesser number of reads uniquely mapped to the pan-transcriptome (63.74%) compared to the Rio reference genome (82.03%), and a higher number of reads mapped to multiple loci in the pan-transcriptome (18.21%) compared to the Rio reference genome (4.4%). In the analysis using the pan-transcriptome, the average number of reads that mapped to too many loci and reads that were unmapped were 1.64% and 16.42% respectively. In the analysis using the Rio reference genome, the average number of reads that mapped to too many loci and reads that were unmapped were 5.08% and 8.5% respectively (Figure 1A). However, despite the overall lower mapping rates in the pan-transcriptome, the total number of assigned hits was higher when compared to the Rio reference genome. In fact, the pan-transcriptome had no unassigned hits while the analysis that used the Rio reference genome had an average of 4.48 million unassigned hits across the samples. (Figure 1B).

**FIGURE 1.**
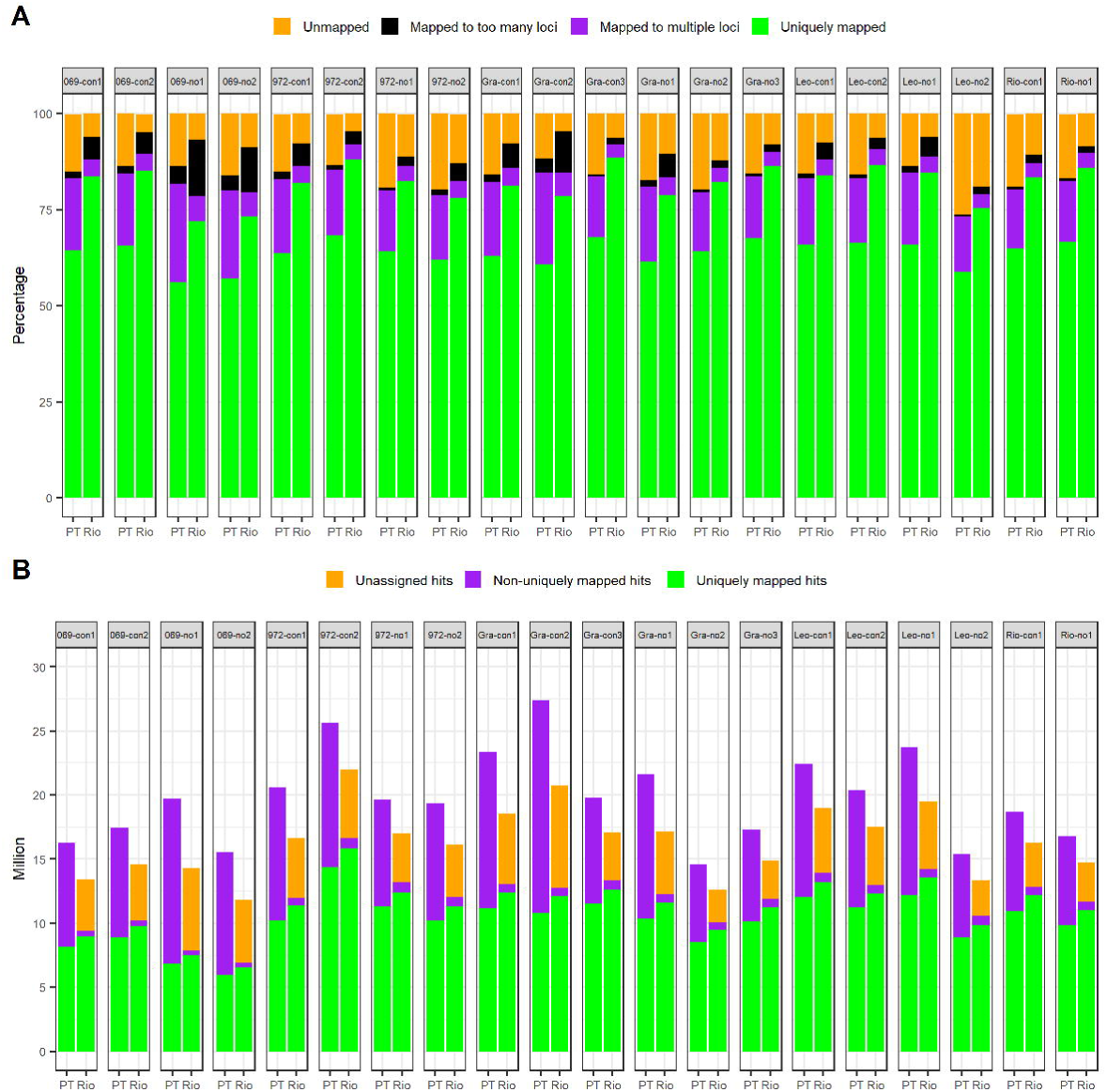
Read alignment and count statistics for the single reference genome analysis (Rio) versus the pan-transcriptome analysis (PT) (A) Alignment metrics from both analyses (in percentages). Green bars indicate uniquely mapped reads that mapped to only one locus. Purple bars show the fraction of reads that mapped to anywhere from 2-10 loci. Black bars represent reads that mapped to more than 10 loci (and were not used in any downstream analyses). Orange bars show the fraction of the reads that did not map to the reference genome/pan-transcriptome; unmapped reads could be as a result of too many mismatches, short read length or no acceptable seeds in reads. (B) Read counts from mmquant (in millions). Green bars indicate uniquely mapped hits that mapped to only one location on the reference genome/pan-transcriptome. Purple bars indicate non-uniquely mapped hits that mapped to more than one location on the reference genome/pan-transcriptome. Orange bars indicate unassigned hits which are hits with no corresponding features on the reference genome/pan-transcriptome. PT, pan-transcriptome; Rio, Rio reference genome.

### 3.2 Single reference genome analysis

#### 3.2.1 Differentially expression analysis

Using a single sweet sorghum genome (Rio) as a reference, 1184, 514 and 76 genes were found to be significantly differentially expressed due to the Treatment, Type and Interaction effects, respectively (Figure 2A). Of these DEGs, 51 were common in all three fixed effects while 60 DEGs were common in Treatment and Type, 14 DEGs were common in Type and Interaction and 5 DEGs were common in Treatment and Interaction. There were 1068, 389 and 6 DEGs unique to Treatment, Type and Interaction effects, respectively.

**FIGURE 2.**
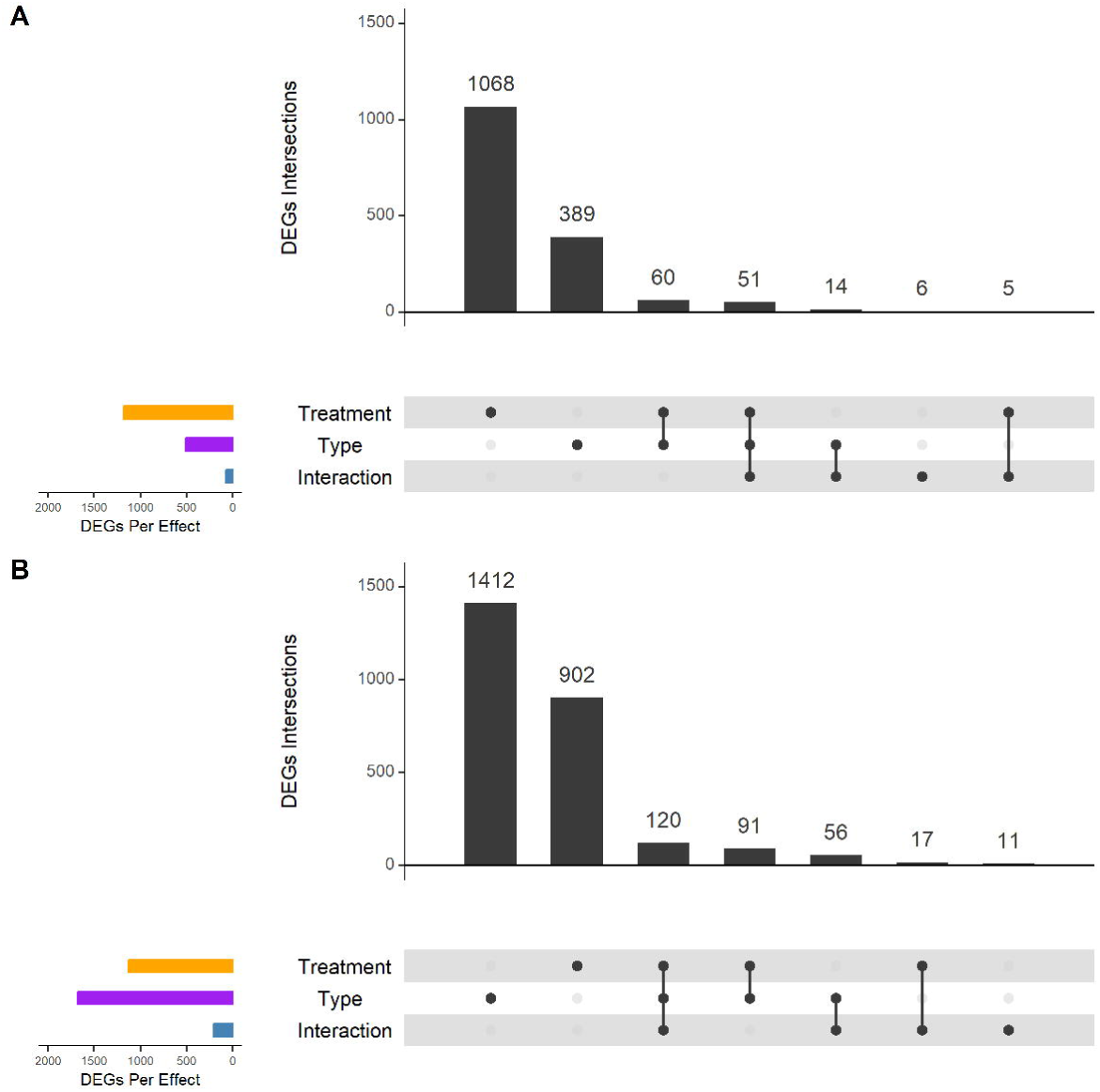
Upset plots show the count of genes expressed due to the model’s fixed effects (Treatment, Type, Interaction) and genes that occur in more than one fixed effect or are unique to a fixed effect. Genes expressed due to the fixed effects are displayed as horizontal bars on the lower left corner of each figure. Treatment, Type and Interaction DEGs are represented by orange, purple and blue bars respectively. DEGs intersection sizes are shown as individual vertical grey bars in each figure with the count of each bar written above it. DEGs that occur in more than one fixed effect are displayed as connected solid black circles under the vertical bars of each figure. Unconnected circles represent DEGs unique to a fixed effect. (A) Analysis using the Rio reference genome. (B) Analysis using the pan-transcriptome. DEG, differentially expressed gene.

#### 3.2.2 Gene ontology enrichment analysis

GO terms enriched among the Treatment DEGs included metal ion binding, copper ion binding, iron ion binding, metal ion transmembrane transporter activity, sucrose synthase activity, heme binding, actin binding, flavin adenine dinucleotide binding and term associated with oxidoreductase activity. Copper ion binding, metal ion binding and iron ion binding share some functional similarities and are close to each other on the semantic space scatterplot. Similarly, terms associated with oxidoreductase activity are close to themselves on the semantic space scatterplot (Figure 3A).

**FIGURE 3.**
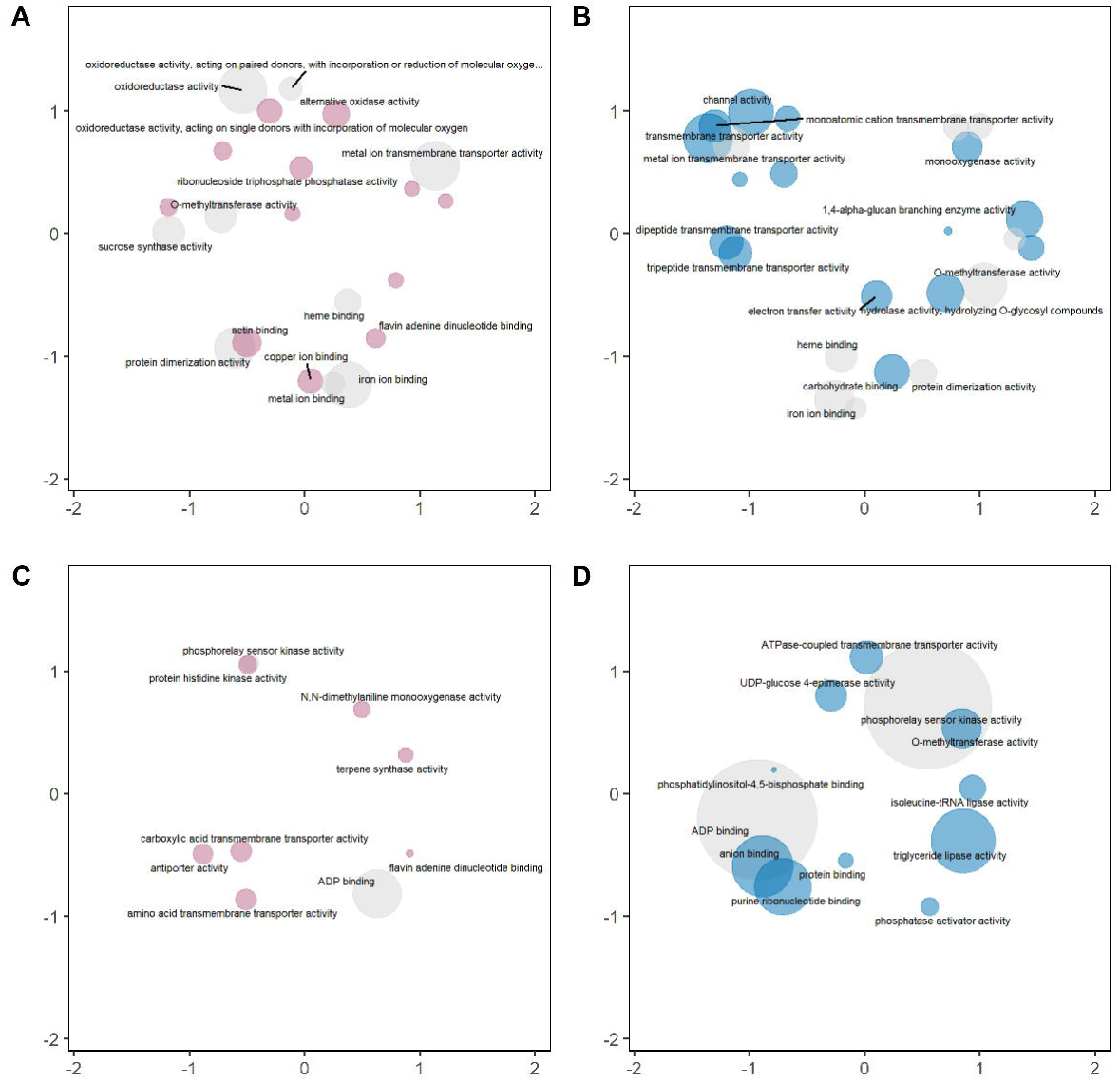
Enriched molecular function GO terms for Treatment and Type DEGs. GO terms are plotted on semantic space x (x-axis) and semantic space y (y-axis). Terms with some functional similarities are close to themselves on the scatterplot. The size of the circle reflects the p-value, with larger circles denoting terms with higher significance. Light-pink colored circles represent terms unique to the Rio reference genome Treatment/Type DEGs, light-blue colored circles represent terms unique to the pan-transcriptome Treatment/Type DEGs and light-grey colored circles represent terms present in both analyses for the fixed effect (Treatment/Type) being considered. (A) Rio reference genome Treatment DEGs (B) Pan-transcriptome Treatment DEGs (C) Rio reference genome Type DEGs (D) Pan-transcriptome Type DEGs DEG, differentially expressed gene; GO, Gene Ontology.

GO terms enriched among the Type DEGs were ADP binding, amino acid transmembrane transporter activity, carboxylic acid transmembrane transporter activity, antiporter activity, phosphorelay sensor kinase activity, protein histidine kinase activity, N, N-dimethylaniline monooxygenase activity, terpene synthase activity and flavin adenine dinucleotide binding (Figure 3C).

GO terms enriched by the Interaction DEGs were oxoglutarate dehydrogenase (succinyl-transferring) activity, chitin binding, serine-type endopeptidase inhibitor activity, N, N-dimethylaniline monooxygenase activity, chitinase activity, O-acyltransferase activity, terpene synthase activity, thiamine pyrophosphate binding (Supplementary Figure 2A).

A full list of GO terms before redundancy reduction can be found in Supplementary File 4.

#### 3.2.3 Enrichment analysis based on evolutionary nodes annotation

More than half (62%) of the genes in the Rio reference genome could be traced back to a particular evolutionary node of origin in the species tree (Figure 4A) based on the orthology analysis. Of the remaining genes, 20 were designated as Single Copy Orthologues, 1295 could not be assigned to any orthogroup, and 12189 could not be unambiguously assigned an age because their copy number pattern did not follow the predicted species tree. For genes that were assigned a node of origin, 836 could be traced to the oldest node in the tree (Plantae), 6617 were common to all Angiosperms, 2516 could be traced to the common ancestor of monocots and dicots (Eudicots), 941 were unique to Monocots, 736 unique to the Poales, 4719 unique to the Poaceae, 1581 unique to the Panicodoideae, and 766 unique to the Andropogoneae. Among the newest gene copies, 935 had an origin in the common ancestor of sorghum and *Miscanthus* (node N15), 971 were common to all sorghums, 674 were common to most, but not all, sorghums (Sorghum subgroup), and 694 were unique to the Rio genotype.

**FIGURE 4.**
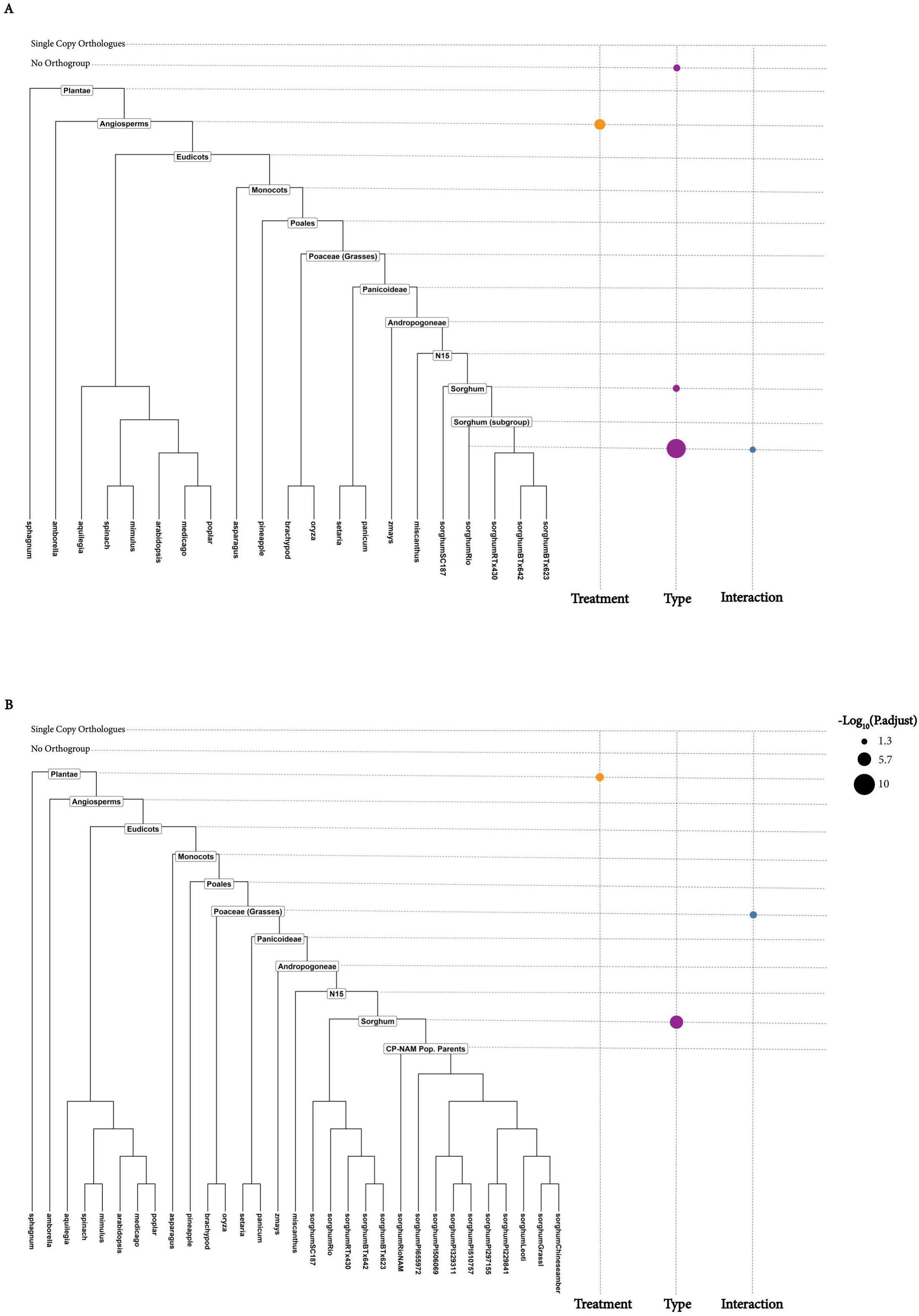
Enrichment analysis based on the age of duplication event. The phylogenetic tree on the left of each figure shows the rooted species tree of the species listed in section 2.6 used in performing orthology analysis with sorghum. Labeled nodes represent the age of duplication events. Genes with duplication events occurring at deeper nodes such as ‘Eudicots’ represent older genes with copies present across all plants that are classified as eudicots while genes with duplication events occurring at shallower nodes such as ‘Sorghum’ represent newer genes with copies present only in the sorghum clade. Categories ‘Single Copy Orthologues’ and ‘No Orthogroup’ represent genes that are single-copy orthologues and those that were unassigned to any orthogroup, respectively. Dot plot to the right of the rooted species tree shows nodes enriched by the Treatment, Type and Interaction DEGs. Orange-colored, purple-colored and blue-colored circles represent Treatment, Type and Interaction DEGs, respectively. Larger circles are more significant (lower p-adjusted values) than smaller circles (higher p-adjusted value). (A) Analysis using the Rio reference genome. (B) Analysis using the pan-transcriptome. DEG, differentially expressed gene

Genes differentially expressed in response to iron stress were significantly enriched for genes that had their origins in the ancestor of all Angiosperms (p=8.58 x 10^-5^). By contrast, the evolutionary origins of genes differentially expressed in response to the difference in genotypes were enriched for the sorghum-specific nodes: Sorghum (p=1.14 x 10^-2^), SorghumRio (p=1.27 x 10^-9^) and No Orthogroups (1.31 x 10^-2^). SorghumRio (p=0.03) was the only evolutionary node enriched among genes differentially expressed due to the interaction effect of iron treatment and genotype (Figure 4A).

### 3.3 Pan-transcriptome

#### 3.3.1 Differentially expressed analysis

There were nearly 70% more genes identified as significantly differentially expressed when aligning to the pan-transcriptome as compared to the analysis performed with a single reference (3013 DEGs with the pan-transcriptome versus 1774 with the Rio reference). The pan-transcriptome analysis uncovered slightly fewer DEGs responding to iron Treatment (1130 vs 1184), but many more DEGs related to Type and Interaction effects (1679 and 204, respectively) (Figure 2B). There were 120 DEGs common in all three fixed effects while 91 DEGs were common in Treatment and Type, 56 DEGs were common in Type and Interaction and 17 DEGs were common in Treatment and Interaction. A total of 902, 1412 and 11 DEGs were unique to Treatment, Type and Interaction effects, respectively.

Comparison of the fixed effects DEGs orthogroups across both analyses showed 579, 318 and 15 orthogroups to overlap for Treatment, Type and Interaction effects, respectively (Figure 5).

**FIGURE 5.**
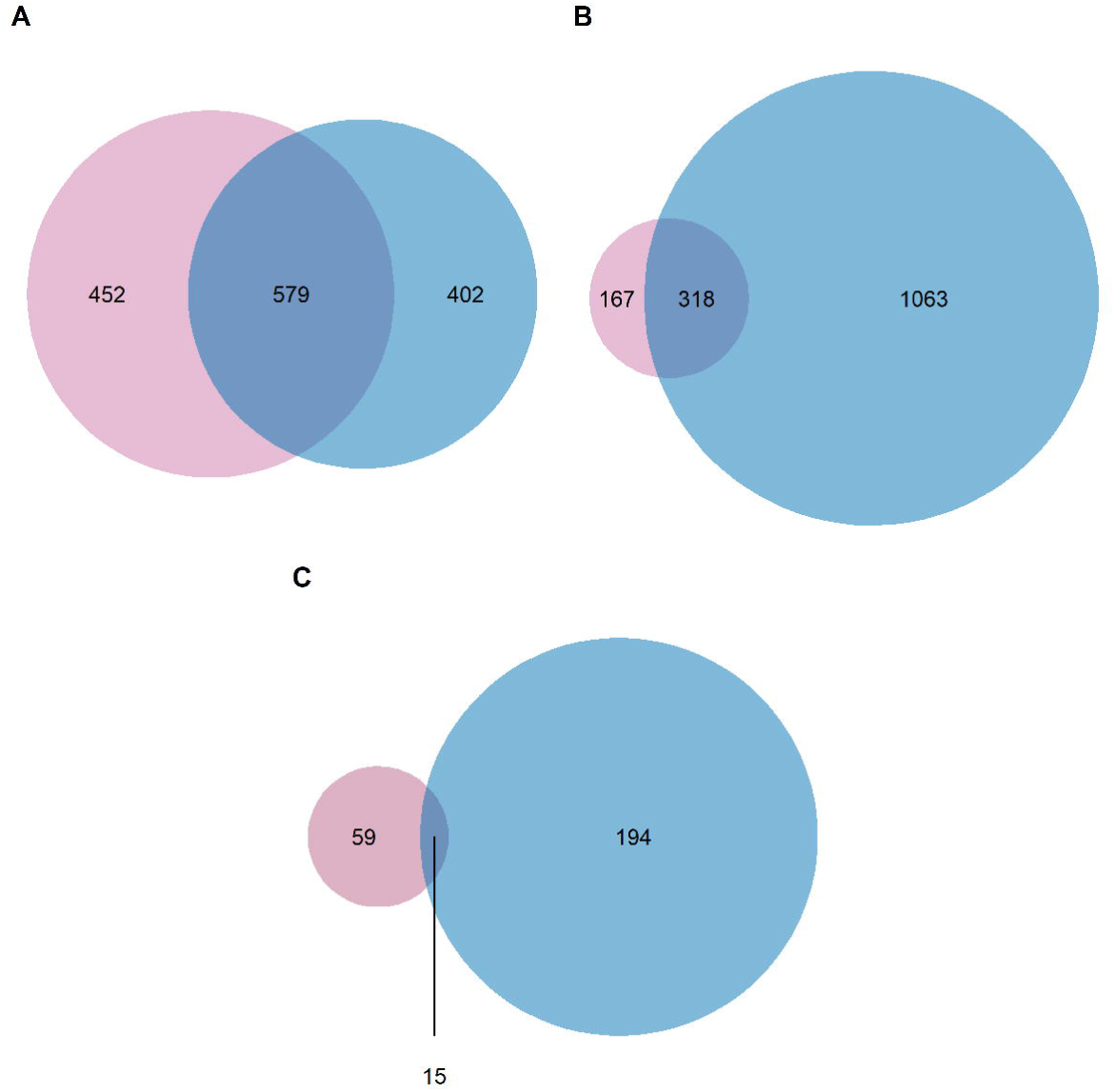
Venn diagram shows DEGs orthogroups that overlap between the analysis that used the Rio reference genome and the analysis that used the pan-transcriptome for each of the model’s fixed effects. The light-pink color represents DEGs orthogroups from the analysis that used the Rio reference genome while the light-blue color represents DEGs orthogroups from the analysis that used the pan-transcriptome. (A) Treatment (B) Type (C) Interaction. DEG, differentially expressed gene

#### 3.3.2 Gene ontology enrichment analysis

Iron ion binding, metal ion binding, and metal ion transmembrane transport activity GO terms enriched among the Treatment DEGs from the analysis that used the Rio reference genome were also enriched among the Treatment DEGs from the analysis that used the pan-transcriptome. Other GO terms enriched among the Treatment DEGs of both analyses were sucrose synthase activity, O-methyltransferase activity, protein dimerization activity, heme binding, and terms associated with oxidoreductase activity (Figures 3A & 3B).

ADP binding and phosphorelay sensor kinase activity GO terms enriched among the Type DEGs from the analysis that used the Rio reference genome were also enriched within the Type DEGs from the analysis that used the pan-transcriptome (Figures 3C & 3D). Other GO terms enriched by pan-transcriptome Type DEGs were triglyceride lipase activity, anion binding, purine ribonucleotide binding, O-methyltransferase activity, ATPase-coupled transmembrane transporter activity, UDP-glucose 4-epimerase activity, isoleucine-tRNA ligase activity, phosphatase activator activity, protein binding, phosphatidylinositol-4,5-bisphosphate binding (Figure 3D).

Only one GO term (chitin binding) significantly enriched in the Interaction DEGs from the analysis that used the Rio reference genome was also enriched in the Interaction DEGs from the analysis that used the pan-transcriptome (Supplementary Figure 2). An interesting GO term enriched by the Interaction DEGs from only the pan-transcriptome analysis was monosaccharide transmembrane transporter activity. A closer look at the DEGs that resulted in this GO term enrichment revealed two sugar transport proteins (SORBI_3002G073600 and SORBI_3001G454100). Other GO terms enriched by pan-transcriptome Interaction DEGs were ADP binding, leucine-tRNA ligase activity, arylformamidase activity, RNA polymerase core enzyme binding, intramolecular transferase activity, and binding (Supplementary Figure 2B).

A full list of GO terms before redundancy reduction can be found in Supplementary File 4.

#### 3.3.3 Enrichment analysis based on evolutionary nodes annotation

The orthology analysis and gene age classification for the genes in the pan-transcriptome resulted in the following evolutionary node assignments: Plantae – 2112, Angiosperms – 9199, Eudicots – 4176, Monocots – 1330, Poales – 843, Poaceae – 5721, Panicoideae – 2098, Andropogoneae – 1274, N15 (sorghum split from *Miscanthus*) – 1511, Sorghum – 3112, CP-NAM Population Parents – 2347, Single Copy Orthologues – 4, No Orthogroup – 2265. 26052 genes could not be unambiguously annotated to any category. Similar to the analysis that used the Rio reference genome, genes found to be significantly differentially expressed in response to iron stress were enriched for a deeper node of origin (Plantae (p=0.003)) while genes that were differentially expressed among genotypes were enriched newer, sorghum-specific gene copies (Sorghum (p=3.2 x 10^-6^))(Figure 4B).

Genes differentially expressed due to the interaction effect of iron treatment and genotype were enriched among genes found in all of the Poaceae (p=0.008) (Figure 4B). This was unlike the analysis that used the Rio reference genome where it was a shallower node that was enriched (SorghumRio) (Figures 4A and 4B).

## 4 Discussion

This study revealed more than a thousand genes significantly differentially expressed in sorghum in response to iron deficiency, many with GO terms relating to iron binding, metal binding, and metal transport. Perhaps more interestingly, this study also uncovered more than 200 genes that responded differently to iron stress in different sorghum types, but the majority of these interaction effects were only apparent when utilizing the pan-transcriptome. This result highlights the benefits of also considering newer, non-core genes in genomics analyses. Gene duplication events are known to play a key role in the adaptation of plants (Kondrashov, 2012; Panchy et al., 2016), and in this study these younger gene copies appear to be the primary drivers of type-specific responses to iron deficiency among these diverse sorghum genotypes.

Iron is integral to plants; so, its deficiency can have a widespread effect that induces other responses in the plant in pathways not immediately connected to metal metabolism. One of the primary motivations for this study was the further elucidation of the relationship between genomic variation in iron transporters and differences in stem sugar accumulation that have been previously reported in sorghum. Studies in other plant species have also found connections between sugar and iron. For example, Zargar et al. (2015b) reported that iron deficiency induces the expression of sugar transporter 4 (*STP4*; AT3G19930) and sugar transporter 13 (*STP13*; AT5G26340). Zargar et al. (2015a) went on to postulate that the high expression of sugar transporters due to iron deficiency might be to increase sugar levels in the shoot for the plant’s fundamental processes. They made this suggestion because iron deficiency leads to a decrease in photosynthetic activity which subsequently leads to a decrease in sugar synthesis. And *STP13* has also been documented to play a role in the reabsorption of sugars from roots under abiotic stress (Yamada et al., 2011). In the present study, a homolog of *STP13* in sorghum (SORBI_3001G454100) was identified as differentially expressed due to iron deficiency in the analysis using the pan-transcriptome. Another sugar transporter (sugar transport protein 8: SORBI_3002G073600) that shares the same conserved domain as SORBI_3001G454100 was also found to be differentially expressed due to iron treatment in the analysis that used the pan-transcriptome. Interestingly, SORBI_3001G454100 and SORBI_3002G073600 both showed significant interaction effects as well as treatment effects, and SORBI_3001G454100 was differentially expressed due to the Type effect as well. Sugar transport proteins 8 and 13 were not identified in any of the analyses that used only the Rio reference genome.

Another interesting group of genes found to be differentially expressed in this study were members of the <u>S</u>ugars <u>W</u>ill <u>E</u>ventually be <u>E</u>xported <u>T</u>ransporters (*SWEET*) gene family. The *SWEET* genes transport sugar across cell membranes and along a concentration gradient (Frank Baker et al., 2012). Members of this gene family have been documented to be potentially responsible for sugar accumulation in the stem of the sweet sorghum genotypes (Mizuno et al., 2016). The analysis that used the pan-transcriptome uncovered significant differences in expression in the following *SWEET* genes: *SWEET1a*, *SWEET1b*, and *SWEET12* in the Treatment group; *SWEET1b* in the Type group; and SWEET1b in the Interaction group. Only *SWEET1a* was identified as differentially expressed in the analysis that used the Rio reference genome and this was only in the Treatment group. Cooper et al. (2019) previously noted that *SWEET1b* (known as *SWEET9_2* in their paper) was differentially expressed between a sweet (Rio) and non-sweet (PR22) sorghum genotype during the soft dough stage, when sweet and non-sweet sorghums exhibit some of the most significant differences in stem sugar content.

Iron deficiency can also impact the expression of genes involved in sucrose synthesis as well as sucrose transport, and levels of sucrose have been noted to be important in regulating iron deficiency response in plant roots (Lin et al., 2016; Chen et al., 2018; Guo et al., 2020). Consistent with this idea, in this study, the GO term GO:0016157 (sucrose synthase activity) was enriched among the Treatment DEGs in both the analysis that used the Rio reference genome and the analysis that used the pan-transcriptome.

Finally, there are a number of other types of genes outside of known sugar metabolism pathways that have also been reported to be responsible for sugar accumulation in the sweet genotypes. Li et al. (2019) compared the expression dynamic of carbon metabolic genes in the internodes of sweet and non-sweet genotypes and reported genes involved in the following process to be differentially expressed: cellulose and monolignol synthesis (*CesA*: cellulose synthase, *PTAL*: L-Phe/-Tyr L-Phe/-Tyr ammonia-lyase and *CCR*: cinnamyl-CoA reductase), starch metabolism (*AGPase*: ADP-glucose pyrophosphorylase, *SS*: soluble starch synthase, *SBE*: starch branching enzyme, and *GPT2*: glucose-6-phosphate-translocator 2), and sucrose metabolism and transport (*TPP*: trehalose-6-phosphate phosphatase, *TPS*: trehalose-6-phosphate synthase and *TST2*: Tonoplast Sugar Transporter). In this study, *PTAL, AGPase, SS, TPP* and *TPS* were found to be differentially expressed due to the genotype effect in the pan-transcriptome analysis but only *AGPase* and *SS* were identified in the analysis that used the Rio reference genome.

## 5 Conclusion

This study revealed that when a pan-transcriptome is used, more than twice as many genes can be identified as differentially expressed due to genotype effect and the interaction effect of genotype and iron treatment. This showcases the utility of the pan-transcriptome in uncovering genes that would have otherwise remained hidden if only a single reference genome was used, and in this case the identification of these additional genes helped to shed light on the processes linking iron metabolism to differences in sugar accumulation in different sorghum types. Newer gene copies were the primary source of differences between the sorghum genotypes while genes expressed as a result of iron stress conditions tended to be older and more evolutionarily conserved, highlighting the utility of the pan-transcriptome in identifying novel targets for crop improvement in the non-core genome a single reference genome might be lacking.

## Data availability statement

The datasets presented in this study can be found on NCBI using the following BioProject accession number: <u>PRJNA1143049</u>. All custom codes used in this study are available at the author’s GitHub repository: https://github.com/Chinaza11/pantranscriptome_iron-deficiency_sorghum-genotypes/

## Author contributions

CDN: Formal analysis, Methodology, Visualization, Writing – original draft, Writing – review & editing. KS: Investigation, Methodology, Writing – review & editing. AAZ: Investigation, Writing – review & editing. WGV: Resources, Writing – review & editing. EAC: Conceptualization, Funding acquisition, Formal analysis, Methodology, Project administration, Supervision, Writing – review & editing

## Acknowledgments

The authors would like to thank Kapeel Chougule and Doreen Ware for their assistance with creating the PAV file used in constructing the pan-transcriptome. The authors would also like to thank Bailee Ku and Tuvani Ramnarine for their assistance with growing plants and performing RNA extractions, and the University Research Computing unit at UNC Charlotte for providing computing resources.

## Publisher’s note

All claims expressed in this article are solely those of the authors and do not necessarily represent those of their affiliated organizations, or those of the publisher, the editors and the reviewers. Any product that may be evaluated in this article, or claim that may be made by its manufacturer, is not guaranteed or endorsed by the publisher.

## Supplementary material

The Supplementary Material for this article can be found online at: https://doi.org/10.5281/zenodo.13900152

## Abbreviations

CP-NAM: Carbon Partitioning Nested Association Mapping
DEG: Differentially Expressed Gene
GO: Gene Ontology

