## Supplementary File for "Utilizing a pan-transcriptome reveals genotype-specific responses to iron deficiency in *Sorghum bicolor*"

### *Supplementary Material*

#### 1 Permutation analysis

Grassl, Leoti and Rio_NAM are classified as sweet genotypes, and PI 506069, PI 655972, and PI297155 are classified as non-sweet genotypes (Voelker et al., 2023). To double-check the result from this original grouping to be sure it is not by chance, a permutation analysis could be carried out. This will involve randomly assigning the genotypes into sweet and non-sweet groups. For example, a group generated by randomly assigning the genotypes to sweet and non-sweet types would be: sweet genotypes (PI 506069, PI 655972, Grassl) and non-sweet genotypes (Leoti, Rio_NAM, PI297155). After randomly assigning the genotypes into groups, differential expression analysis is conducted on each of the groups. Each gene that was differentially expressed in the original grouping is investigated in the differential expression analysis result from the random groupings to see the number of times they are also differentially expressed and finally, a permutation p-value for this gene is computed. Genes with permutation p-values less than or equal to 0.05 are said to be significant after the permutation test.

Ideally, a minimum of 500 random groupings is needed to ensure this permutation test but the limitation of this study is the only possible number of random groupings is 20. We still went ahead and did a permutation analysis though its result can't be entirely accepted, and Supplementary Figures 3-6 and Supplementary File 4 show these results.

After the permutation test, in the analysis that used the Rio reference genome, 227 out of the 514 Type DEGs and 32 out of the 76 Interaction DEGs were reported to not have been significantly present in the random groupings. In the analysis that used the pan-transcriptome, 657 out of the 1679 Type DEGs and 101 out of the 204 Interaction DEGs were reported to not have been significantly present in the random groupings.

All other results are largely similar to the results in the main paper and the same conclusions in the main paper can also be drawn from the results obtained after the permutation test.

### 2 Supplementary Figures and Tables

#### 2.1 Supplementary Figures

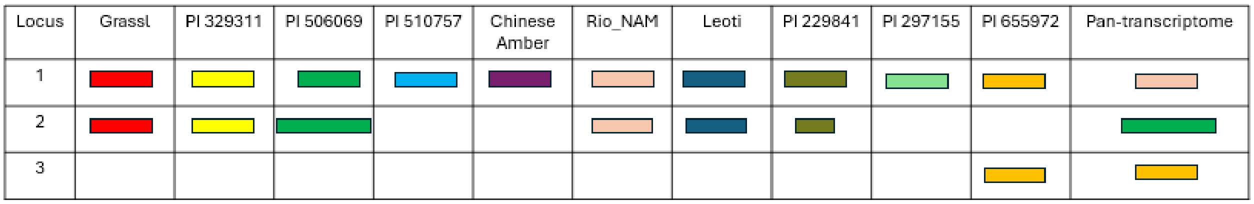


SUPPLEMENTARY FIGURE 1

Pan-transcriptome construction. The longest sequence for each locus was selected to be the representative sequence. If Rio_NAM also had the longest sequence, it was picked over the other founder parent.

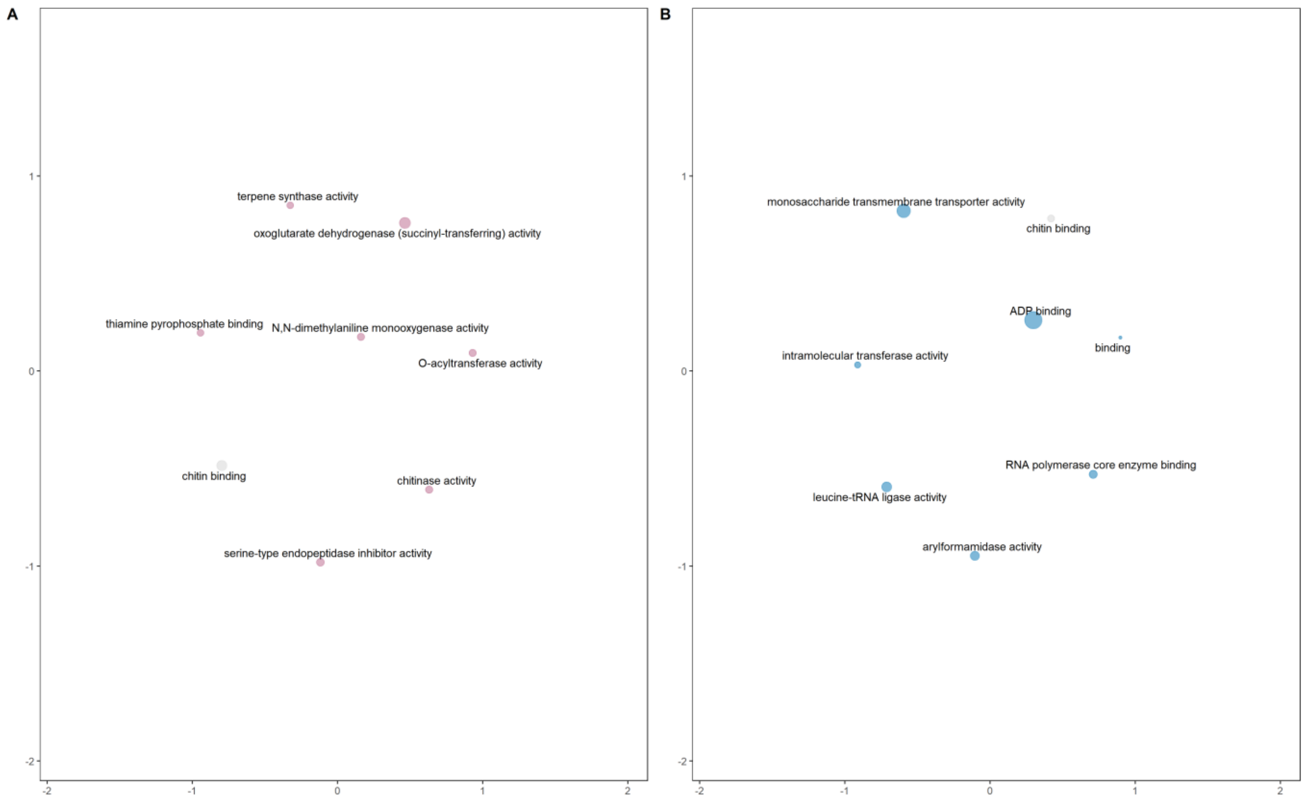


SUPPLEMENTARY FIGURE 2

Enriched molecular function GO terms for Interaction DEGs. GO terms are plotted on semantic space x (x-axis) and semantic space y (y-axis). Terms with some functional similarities are close to themselves on the scatterplot. The size of the circle reflects the p-value, with larger circles denoting terms with higher significance. Light-pink colored circles represent terms unique to the Rio reference genome DEGs, light-blue colored circles represent terms unique to the pan-transcriptome DEGs and light-grey colored circles represent terms present in both analyses (A) Rio reference genome Interaction DEGs (B) Pan-transcriptome Interaction DEGs

DEG, differentially expressed gene; GO, Gene Ontology.

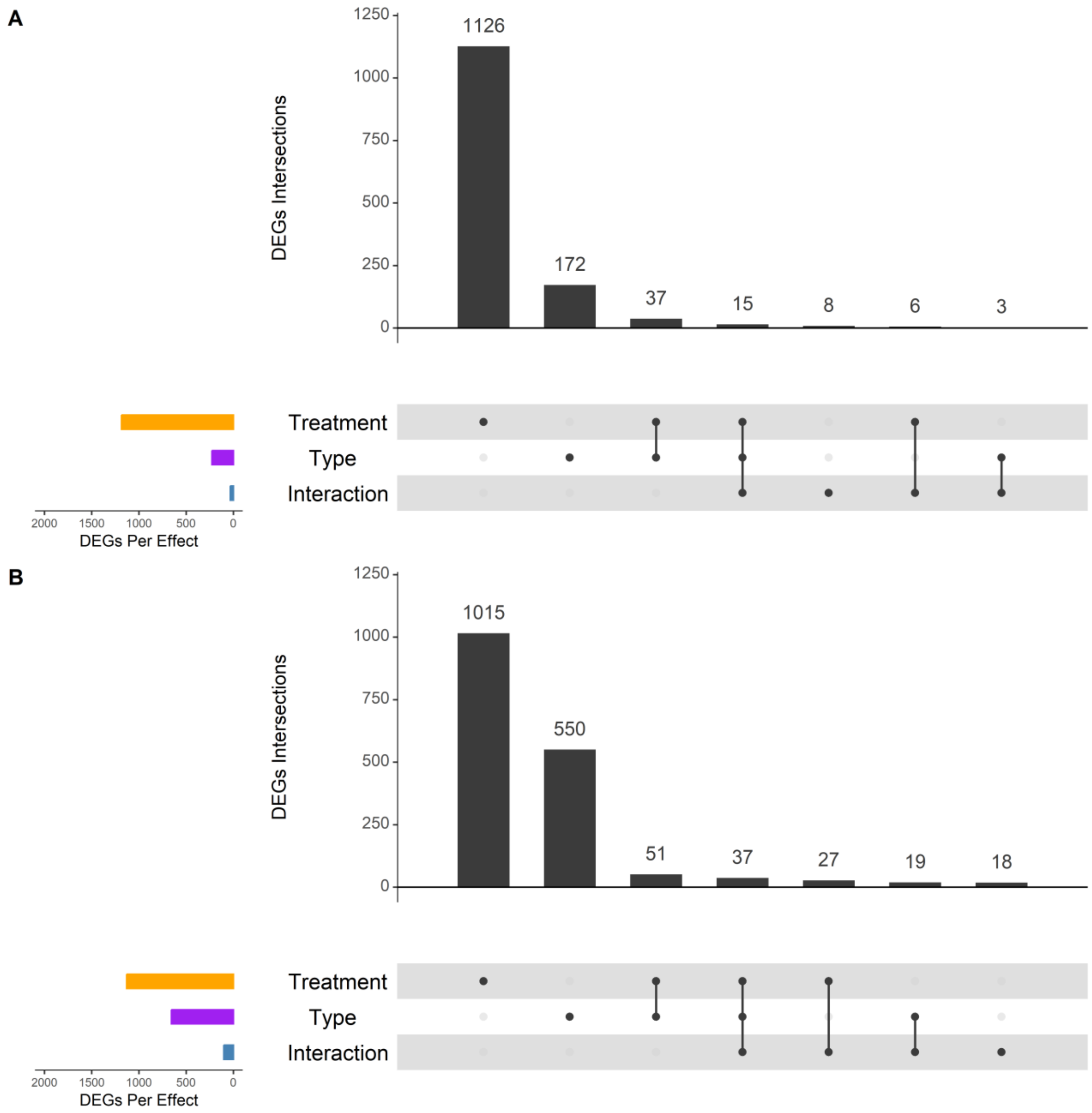


SUPPLEMENTARY FIGURE 3

Upset plots show the count of genes after the permutation test that are expressed due to the model's fixed effects (Treatment, Type, Interaction) and that occur in more than one fixed effect or are unique to a fixed effect. Genes expressed due to the fixed effects are displayed as horizontal bars on the lower left corner of each figure. Treatment, Type and Interaction DEGs are represented by orange, purple and blue bars respectively. DEGs intersection sizes are shown as individual vertical grey bars in each figure with the count of each bar written above it. DEGs that occur in more than one fixed effect are displayed as connected solid black circles under the vertical bars of each figure. Unconnected circles represent DEGs unique to a fixed effect. (A) Analysis using the Rio reference genome. (B) Analysis using the pan-transcriptome.

DEG, differentially expressed gene.

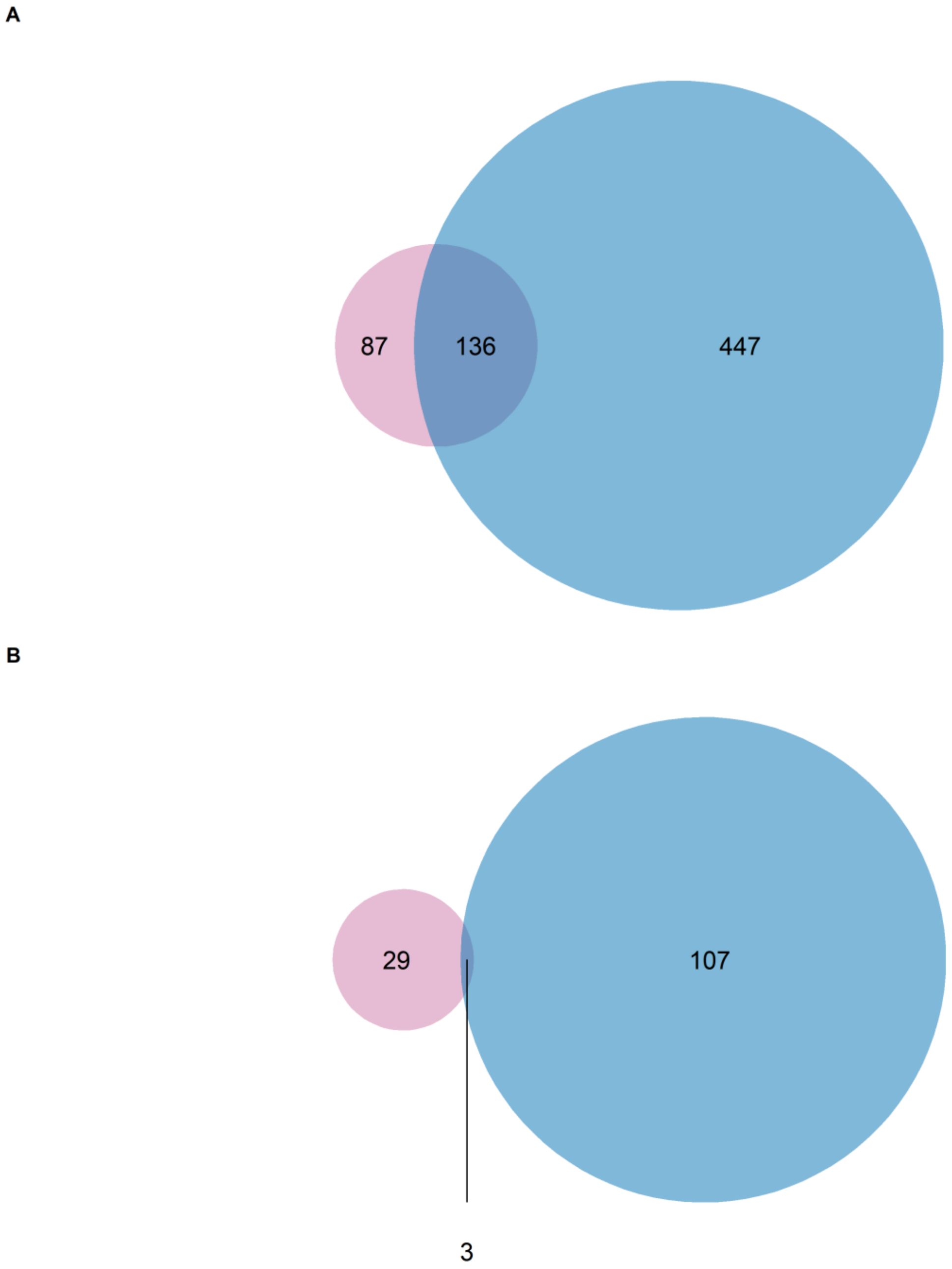


SUPPLEMENTARY FIGURE 4

Venn diagram shows DEGs orthogroups that overlap between the analysis that used the Rio reference genome and the analysis that used the pan-transcriptome for each of the model's fixed effects. The light-pink color represents DEGs orthogroups from the analysis that used the Rio reference genome while the light-blue color represents DEGs orthogroups from the analysis that used the pan-transcriptome. (A) Type (B) Interaction.

DEG, differentially expressed gene

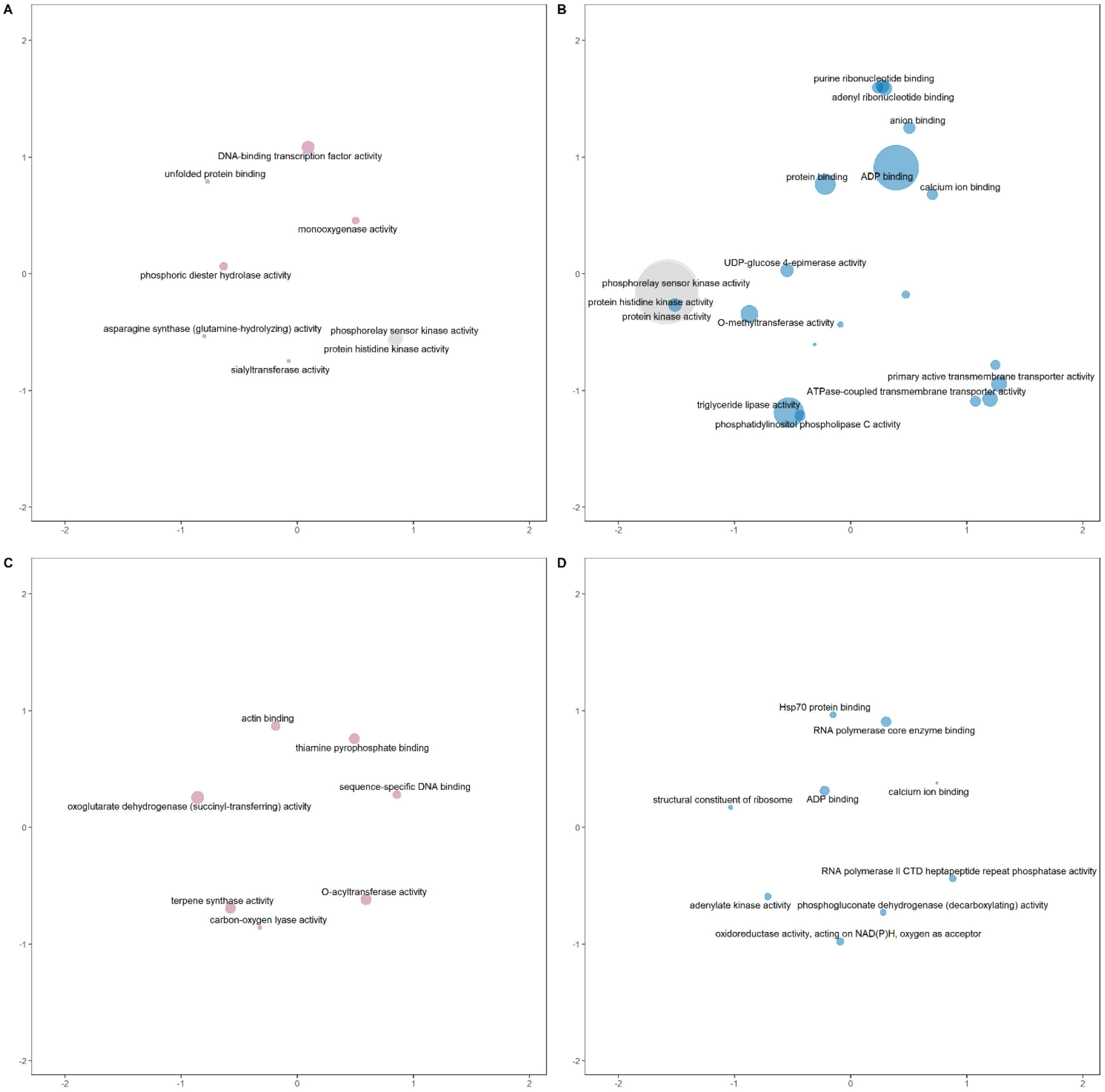


SUPPLEMENTARY FIGURE 5

Enriched molecular function GO terms for Type and Interaction DEGs. GO terms are plotted on semantic space x (x-axis) and semantic space y (y-axis). Terms with some functional similarities are close to themselves on the scatterplot. The size of the circle reflects the p-value, with larger circles denoting terms with higher significance. Light-pink colored circles represent terms unique to the Rio reference genome Type/Interaction DEGs, light-blue colored circles represent terms unique to the pan-transcriptome Type/Interaction DEGs and light-grey colored circles represent terms present in both analyses for the fixed effect (Type/Interaction) being considered. (A) Rio reference genome Type DEGs (B) Pan-transcriptome Type DEGs (C) Rio reference genome Interaction DEGs (D) Pan-transcriptome Interaction DEGs

DEG, differentially expressed gene; GO, Gene Ontology.

A

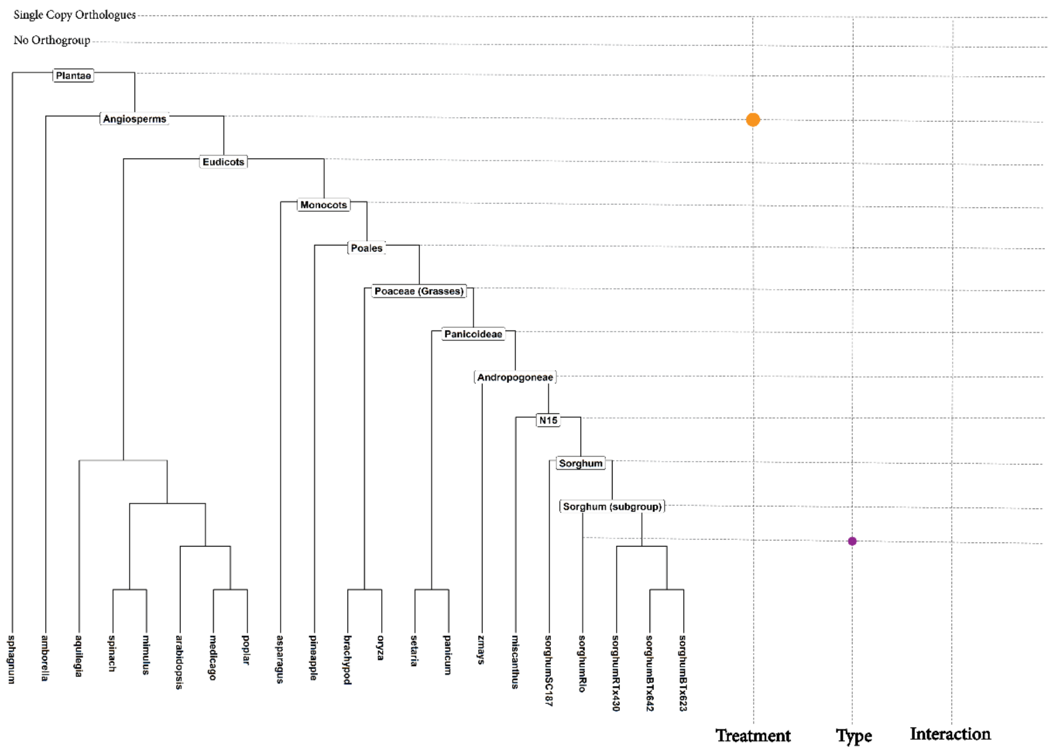
Single Copy Orthologues
No Orthogroup
Plantae
Angiosperms
Eudicots
Monocots
Poales
Poaceae (Grasses)
Panicoideae
Andropogoneae
N15
Sorghum
Sorghum (subgroup)
sphagnum
amborella
aquilegia
spinach
mimulus
arabidopsis
medicago
poplar
asparagus
pineapple
brachypod
oryza
setaria
panicum
zmays
miscanthus
sorghumSC187
sorghumRio
sorghumRTx430
sorghumBTx642
sorghumBTx623
Treatment
Type
Interaction

B

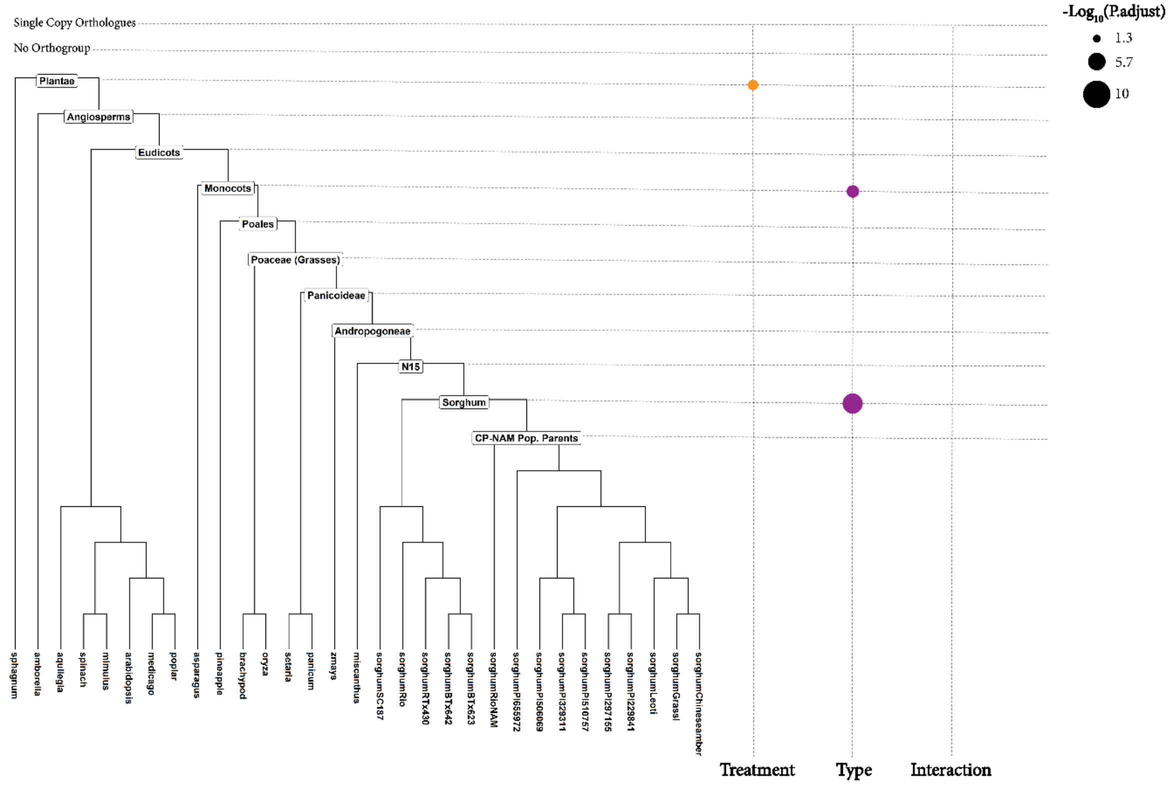
-Log10(P.adjust)
1.3
5.7
10
Single Copy Orthologues
No Orthogroup
Plantae
Angiosperms
Eudicots
Monocots
Poales
Poaceae (Grasses)
Panicoideae
Andropogoneae
N15
Sorghum
CP-NAM Pop. Parents
sphagnum
amborella
aquilegia
spinach
mimulus
arabidopsis
medicago
poplar
asparagus
pineapple
brachypod
oryza
setaria
panicum
zmays
miscanthus
sorghumSC187
sorghumRio
sorghumRTx430
sorghumBTx642
sorghumBTx623
sorghumRioNAM
sorghumPI655972
sorghumPI506069
sorghumPI329311
sorghumPI510757
sorghumPI297155
sorghumPI229841
sorghumLeoti
sorghumGrassl
sorghumChineseamber
Treatment
Type
Interaction

SUPPLEMENTARY FIGURE 6

Enrichment analysis based on the age of duplication event. The phylogenetic tree on the left of each figure shows the rooted species tree of the species listed in section 2.6 used in performing orthology analysis with sorghum. Labeled nodes represent the age of duplication events. Genes with duplication events occurring at deeper nodes such as ‘Eudicots’ represent older genes with copies present across all plants that are classified as eudicots while genes with duplication events occurring at shallower nodes such as ‘Sorghum’ represent newer genes with copies present only in the sorghum clade. Categories ‘Single Copy Orthologues’ and ‘No Orthogroup’ represent genes that are single-copy orthologues and those that were unassigned to any orthogroup, respectively. Dot plot to the right of the rooted species tree shows nodes enriched by the Treatment, Type and Interaction DEGs. Orange-colored, purple-colored and blue-colored circles represent Treatment, Type and Interaction DEGs, respectively. Larger circles are more significant (lower p-adjusted values) than smaller circles (higher p-adjusted value). A) Analysis using the Rio reference genome. (B) Analysis using the pan-transcriptome.

DEG, differentially expressed gene

#### 2.2 Supplementary Tables

SUPPLEMENTARY TABLE 1
Genotypes and replicates used in this study.

| Genotype name | Condition | Replicate | Sample ID | Type |
|---|---|---|---|---|
| Grassl | Control (100 µM Fe) | 1 | Gra-con1 | Sweet |
| | | 2 | Gra-con2 | |
| | | 3 | Gra-con3 | |
| | Low iron (0 µM Fe) | 1 | Gra-no1 | |
| | | 2 | Gra-no2 | |
| | | 3 | Gra-no3 | |
| Leoti | Control (100 µM Fe) | 1 | Leo-con1 | |
| | | 2 | Leo-con2 | |
| | Low iron (0 µM Fe) | 1 | Leo-no1 | |
| | | 2 | Leo-no2 | |
| Rio | Control (100 µM Fe) | 1 | Rio-con1 | |
| | Low iron (0 µM Fe) | 1 | Rio-no1 | |
| PI 506069 | Control (100 µM Fe) | 1 | 069-con1 | Non-sweet/Biomass |
| | | 2 | 069-con2 | |
| | Low iron (0 µM Fe) | 1 | 069-no1 | |
| | | 2 | 069-no2 | |
| PI 655972 | Control (100 µM Fe) | 1 | 972-con1 | |
| | | 2 | 972-con2 | |
| | Low iron (0 µM Fe) | 1 | 972-no1 | |
| | | 2 | 972-no2 | |

SUPPLEMENTARY TABLE 2

Surviving reads after trimming (in millions).

| Sample ID | Read | Sample ID | Read |
|---|---|---|---|
| 069-con1 | 12.7 | Gra-con3 | 17 |
| 069-con2 | 13.5 | Gra-no1 | 16.9 |
| 069-no1 | 12.2 | Gra-no2 | 13.4 |
| 069-no2 | 10.5 | Gra-no3 | 15 |
| 972-con1 | 16 | Leo-con1 | 18.4 |
| 972-con2 | 21 | Leo-con2 | 17 |
| 972-no1 | 17.6 | Leo-no1 | 18.5 |
| 972-no2 | 16.5 | Leo-no2 | 15.1 |
| Gra-con1 | 17.7 | Rio-con1 | 17 |
| Gra-con2 | 17.8 | Rio-no1 | 14.9 |
